# Nucleocytoplasmic transport senses mechanics independently of cell density in cell monolayers

**DOI:** 10.1101/2024.01.11.575167

**Authors:** Ignasi Granero-Moya, Guillaume Belthier, Bart Groenen, Marc Molina-Jordán, Miguel González-Martín, Xavier Trepat, Jacco van Rheenen, Ion Andreu, Pere Roca-Cusachs

## Abstract

Cells sense and respond to mechanical forces through mechanotransduction, which regulates processes in health and disease. In single cells, mechanotransduction involves the transmission of force to the cell nucleus, where it affects nucleocytoplasmic transport (NCT) and the subsequent nuclear localization of transcriptional regulators such as YAP. However, if and how NCT is mechanosensitive in multicellular systems is unclear. Here, we characterize and use a fluorescent sensor of nucleocytoplasmic transport (Sencyt) and demonstrate that nucleocytoplasmic transport responds to mechanics but not cell density in cell monolayers. Using monolayers of both epithelial and mesenchymal phenotype, we show that NCT is altered in response both to osmotic shocks, and to the inhibition of cell contractility. Further, NCT correlates with the degree of nuclear deformation measured through nuclear solidity, a shape parameter related to nuclear envelope tension. In contrast and in opposition to YAP, NCT is not affected by cell density, showing that the response of YAP to both mechanics and cell-cell contacts operates through distinct mechanisms. Our results demonstrate the generality of the mechanical regulation of NCT.

## Introduction

Cells sense and respond to their mechanical context in a process called mechanotransduction. Mechanotransduction is essential in physiological situations such as organ development (Hamant & Saunders, 2020) or embryogenesis (Brunet et al., 2013) and also in pathological settings, for instance tumour progression (Broders-Bondon et al., 2018). One of the cell elements involved in mechanotransduction is the cell nucleus, which responds to both intracellular and extracellular forces through several mechanisms. These mechanisms involve changes in chromatin architecture (Nava et al., 2020), in the conformation and localization of nucleoskeletal elements such as lamins (Philip & Dahl, 2008; Swift et al., 2013), in nuclear membrane tension (Lomakin et al., 2020; Venturini et al., 2020), and in the localization and activity of transcriptional regulators (Elosegui-Artola et al., 2017; Tajik et al., 2016). In single cells, transcriptional regulators including YAP (Elosegui-Artola et al., 2017), Twist, Snail, and SMAD3 (Andreu et al., 2022) localize to the nucleus in response to force due to changes in nucleocytoplasmic transport (NCT). Specifically, force applied to the nucleus increases nuclear membrane tension, NPC diameter, and diffusion through NPCs (Elosegui-Artola et al., 2017; Schuller et al., 2021; Zimmerli et al., 2021). Both passive and facilitated diffusion (i.e., passive and active transport) are affected by force, but to different extents. This causes a differential effect that leads to force-dependent nuclear or cytoplasmic accumulation of proteins depending on the balance between their passive transport properties and their active transport properties (governed by their nuclear localization or export sequences) (Andreu et al., 2022).

The role of NCT in mechanotransduction is thus established for single cells, but if and how it applies to multicellular systems is unclear. In multicellular systems, cell mechanotransduction involves a complex interplay between cell-matrix and cell-cell adhesion (Aragona et al., 2013; Maniotis et al., 1997). Further, cell-cell adhesion per se also regulates transcriptional regulators such as YAP, in ways that could be independent of mechanotransduction mechanisms (Aragona et al., 2013; Zhao et al., 2007, 2008). Thus, to what extent NCT changes can explain mechanotransduction responses in multicellular systems is unknown. To address this, we need a NCT reporter which is sensitive to mechanical forces, but not to signaling pathways (such as the Hippo pathway that regulates YAP). In our previous work (Andreu et al., 2022), we screened a battery of synthetic constructs that expressed inert, freely diffusing proteins, that only interact with the active transport machinery through nuclear localization sequences (NLS). These proteins showed different facilitated and passive transport rates, and some of them had mechanosensitive shuttling rates and localization. In single fibroblasts, the synthetic protein L_NLS-41 kDa (Figure 1a) presented the biggest mechanosensitivity, defined as the change in localization in response to force. Indeed, in response to force applied to the nucleus, L_NLS-41 kDa showed increased rates of both passive and active nuclear transport (Figure 1b). However, active transport was more affected by force, leading to a force-dependent accumulation in the nucleus.

**Figure 1.**
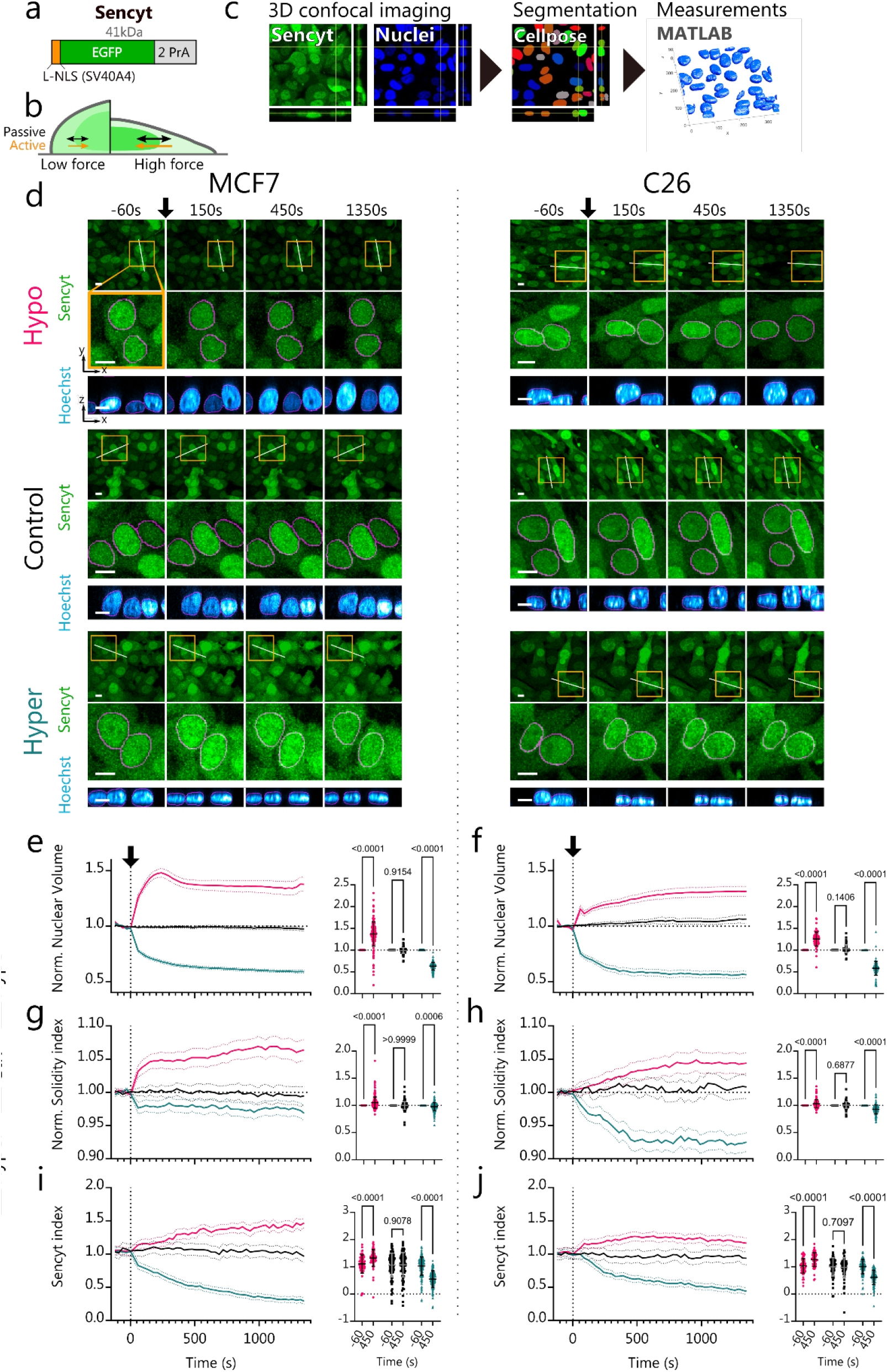
Hypo- and hyper-osmotic shocks increase and decrease nucleocytoplasmic transport respectively. a) schematic representation of Sencyt. Its elements include i) an NLS (SV40A4) based on that from the SV40 virus, but with a point mutation to reduce affinity to importins. ii) An EGFP molecule for visualization. iii) 2 repeats of the inert protein PrA to confer a final molecular weight of 41 kDA, thereby regulating passive diffusion. b) Scheme of Sencyt response to force. When force exerted to the nucleus increases, passive diffusion increases, but active transport increases even more. This increases Sencyt nuclear localization (Andreu et al., 2022). c) Image processing workflow chart. Cells stably transfected with Sencyt are imaged for Sencyt and nuclei (Hoechst label). Nuclei are then segmented, and nuclei shape parameters and Sencyt index are calculated (see methods). d) representative images as a function of time of cells submitted to hypo-, control, or hyper-osmotic shocks, both for MCF7 and C26 cell lines. Arrow indicates the beginning of treatment. In top panels, yellow squares indicate zoomed areas in middle panel, white lines indicate the location of vertical nuclear cross-sections shown in bottom panel. Magenta lines show nuclear mask limits. Scale bar is 10 µm. e,f) Nuclear volume measurements normalized to the 5 first timepoints, and statistics pre/post treatment (N=170, 130, 188, 81, 72, 107 cells) f,h) Solidity index measurements normalized to the 5 first timepoints and statistics pre/post treatment (N=230, 170, 231, 121, 104, 142 cells) i,j) Sencyt index measurements and statistics pre/post treatment (N=66, 101, 152, 60, 63, 103 cells). Sencyt index is defined as the logarithm in base 2 of the ratio of the mean nuclear fluorescence and the mean cytoplasmic fluorescence (see Methods). p-values calculated with 2-way ANOVA corrected with Šídák’s multiple comparisons test. Error bars represent 95% CI for timelapse graphs and SD for statistical graphs. All data include 3 independent repeats.

Due to these properties, L_NLS-41 kDa is an appropriate mechanosensitive NCT reporter, which for convenience we have renamed as **SE**nsor of **N**ucleo**CY**toplasmic **T**ransport (Sencyt). In this work, we use Sencyt in cell monolayers, and show that NCT responds to mechanical inputs but not cell-cell contacts, thereby separating the two types of inputs that are the major regulators of YAP.

## RESULTS

To evaluate the role of NCT in the mechanotransduction of multicellular systems, we used 2 different cell lines stably expressing Sencyt: MCF7 and C26. Both are cancer cell lines, but present different characteristics. MCF-7 are epithelial cells isolated from metastatic adenocarcinoma of a human breast tumor and are used for breast cancer research and many mechanobiological studies. They have an epithelial phenotype (Ahlstrom & Erickson, 2007), with strong cell-cell adhesions. C-26 is a murine colon adenocarcinoma cell line (also named MCA-26, CT-26, and Colo-26) (Corbett et al., 1975). It has a more mesenchymal phenotype, presenting thus an interesting contrast to MCF7. To test the mechanosensitivity of NCT in these two cell lines in a multicellular context, we carried out two types of mechanical perturbations: osmotic shocks, and inhibition of the forces exerted by the cell actin cytoskeleton.

### Hypo- and hyper-osmotic shocks increase and decrease nucleocytoplasmic transport respectively

Osmotic shocks have been widely used to alter mechanical conditions of the nucleus (Schuller et al., 2021; Venturini et al., 2020; Zimmerli et al., 2021). In this work, we have used the osmotic stress conditions previously used in similar works (Elosegui-Artola et al., 2017), to induce nuclear swelling or shrinking, thereby affecting the tension in the nuclear envelope (Dahl et al., 2004; Enyedi et al., 2016), which in turn affects NPC diameter (Zimmerli et al., 2021). We applied the osmotic shocks on cells while we imaged confocally Sencyt and the nucleus (through Hoechst staining) (Figure 1c). The well-described responses for hypo-osmotic shocks include an inflow of water into the cell that causes an increase of cell and nuclear volumes, and a decrease in the concentration of solutes inside of the cell (Churney, 1942; Finan et al., 2009; Lemière et al., 2022). Opposite to hypo-osmotic shocks, hyper-osmotic shocks cause an outflow of water from the cell, that causes a decrease in the cell and nuclear volumes and increase the concentration of solutes (Churney, 1942; Finan et al., 2009; Lemière et al., 2022). To track changes in nuclear volume, and nuclear shape in general, we segmented nuclear images in 3D, and calculated different shape parameters (see methods and Supp. figure 1). By measuring volume changes, we reproduce these trends (Figure 1d-f and Supp. figure 2). For both cell lines, hypo-osmotic shocks increased nuclear volume (by a 50% for MCF7 and a 30% for C26). Inversely, hyper-osmotic shocks reduced nuclear volume (up to 40% in both cell lines, Figure 1e.-f, Supp. figure 2a-b). In C26, the hypo-osmotic shock changes are milder than in MCF7, potentially due to different properties of the nucleoskeleton or initial differences in cell and nuclear osmolarity, which resists nuclear deformations. In MCF7, we also observed an initial nuclear volume increase by a 50%, followed by a decrease to a 40%. This may be explained by an adaptative mechanism, by which cells decrease hypo-osmotic stress by reducing the internal ion concentration (Enyedi et al., 2016; Hoffmann et al., 2009; Lang et al., 1998).

The nucleus is delimited by a double lipidic membrane that is not elastic (Hallett et al., 1993; Needham & Nunn, 1990). Assuming it is finite, increasing the volume of a wrinkled nucleus should, first, increase the nuclear membrane area until the exhaustion of the membrane reservoirs, and second, decrease the number of wrinkles and make the nucleus smoother by increasing nuclear membrane tension, as previously suggested (Niethammer, 2021). Regarding the first part, nuclear surface area increased/decreased for both cell lines when submitted to hypo/hyper-osmotic shocks (Supp. figure 2e-f). To tackle the second part, we measured the nuclear Solidity index (see methods). The Solidity index quantifies the overall concavity of a 3D volume, with high values corresponding to a taut nucleus, and low values corresponding to a wrinkled nucleus (Supp. figure 1). It can thus be understood as an indirect assessment of nuclear membrane tension, since we would expect a high Solidity index for a nucleus submitted to high membrane tension. Consistent with this framework, the Solidity index increased in the hypo shock condition, and decreased in the hyper shock condition (Figure 1g-h), suggesting changes in nuclear envelope tension.

Then, we measured the Sencyt index, defined as the logarithm (in base 2) of the nuclear-to-cytoplasmic ratio of Sencyt signal (see methods). Thus, a positive Sencyt index indicates nuclear localization, a negative one indicates cytoplasmic localization, and zero an equal distribution between both compartments. By measuring changes in the Sencyt index, we can track the ability of the cell NCT system to localize a cargo protein in the nucleus. Upon hypo-osmotic shocks, the Sencyt index increased along with nuclear volume and solidity (Figure 1i-j). On the other hand, upon hyper-osmotic shocks, the Sencyt index decreased along with nuclear volume and solidity (Figure 1i-j). As a control, we transfected the cells with mCherry, which behaves in a completely diffusive way, and occupies evenly all accessible spaces in the cell. The localization of mCherry was not affected by osmotic shocks, except for a small increase in response to hyper-osmotic shocks in MCF7 cells (i.e., in the opposite direction to the effect on Sencyt, Supp. figure 2g-l)). Thus, the changes in Sencyt index are not due to potential effects in the water fluxes, geometry, available space, or effective volume that Sencyt can occupy in the two compartments. Therefore, our data show that osmotic shocks regulate the ability of NCT to accumulate cargoes in the nucleus, in a manner consistent with a role of nuclear envelope tension.

### Myosin II and Arp2/3 inhibition decrease NCT

As a second mechanical perturbation, we inhibited actomyosin activity. To this end, we combined 25µM para-NitroBlebbistatin (to inhibit Myosin II) and 50µM CK666 (to inhibit Arp2/3 and therefore actin branching) (Képiró et al., 2014; Nolen et al., 2009). Of note, we combined both drugs because, in an epithelial context, Para-NitroBlebbistatin alone is not sufficient to deplete nuclear mechanotransduction. Indeed, myosin contractility inhibition alone can lead to increased cell spreading and nuclear deformation, increasing (rather than decreasing) YAP nuclear concentration (Kechagia et al., 2023).

In this set-up, we performed 3 conditions in parallel: (1) a negative control treated with the vehicle, (2) a positive control treated with the drug combination, and (3) a drug washout condition. In the drug washout condition the drugs were washed out after 2 h of imaging (Figure 2a). Then, we analyzed the changes in Sencyt index with time in all three conditions. Treating the cells with the drug combination decreased the Sencyt index when compared with non-treated cells (Figure 2b-c). In the case of the drug washout condition, retrieving the treatment restored the levels of NCT after 1h of drug washout, though not entirely (Figure 2b-c). Surprisingly, when we checked for changes in nuclear volume and Solidity index before and after the treatment there were no remarkable trends and no significant changes (Supp. figure 3a-d). Thus, mechanical force can affect NCT, even without clear nuclear deformations. To explain this, we hypothesize that there may be changes in nuclear envelope tension without requiring high deformations, as the nuclear envelope is a planar mechanical stiff material (Hallett et al., 1993; Needham & Nunn, 1990).

**Figure 2.**
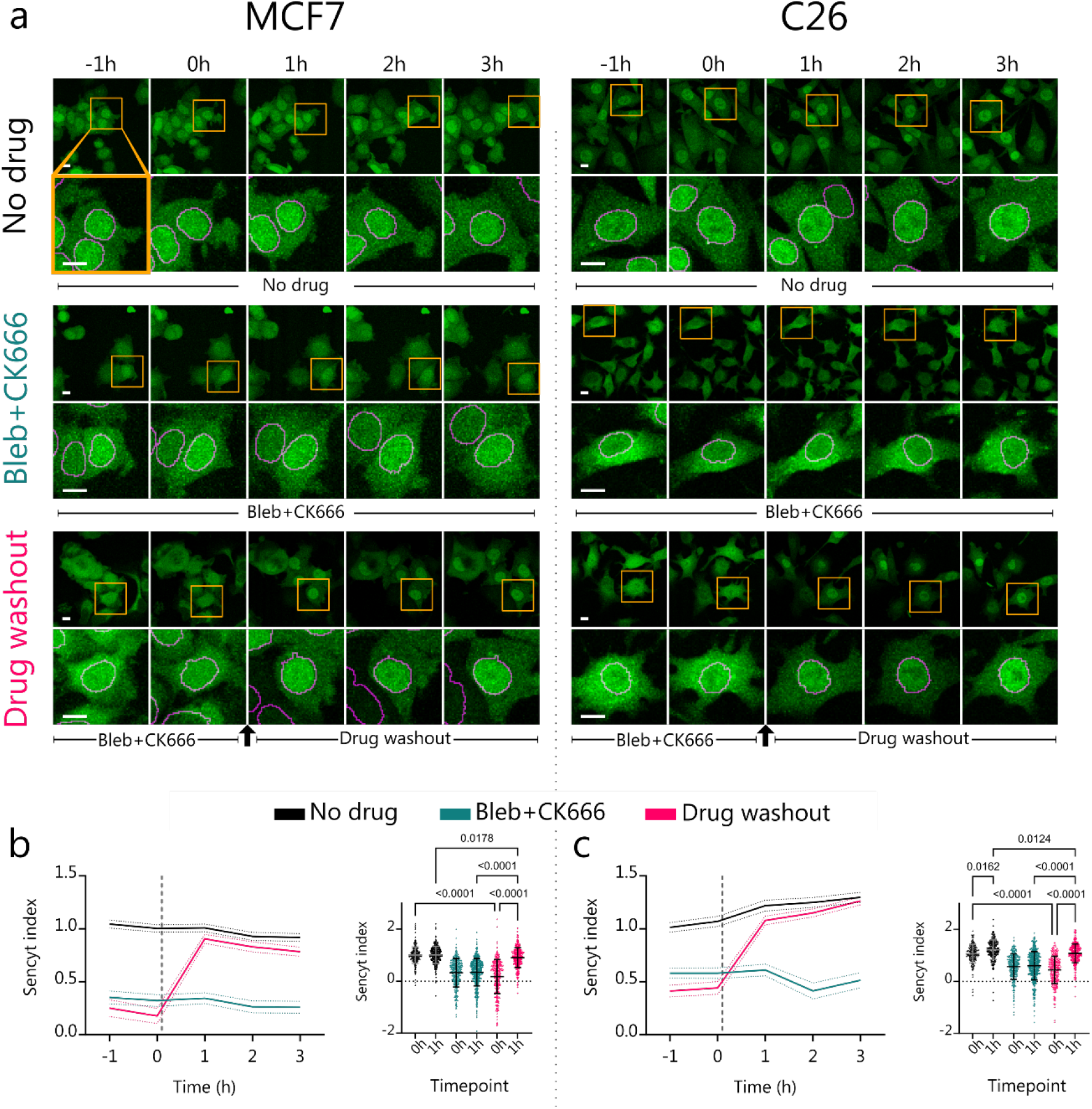
Myosin II and Arp2/3 inhibition decreases NCT. a) representative images as a function of time of cells submitted to different treatments, both for MCF7 and C26 cell lines. In top panels, yellow squares indicate zoomed areas in middle panel. Magenta lines show nuclear mask limits. Scale bar is 10 µm. b,c) Corresponding measurements and statistics for Sencyt index. (N=301, 307, 376, 413, 327, 299 cells). p-values calculated with Kruskal-Wallis test corrected with Dunn’s multiple comparisons test. Error bars represent 95% CI for timelapse graphs and SD for statistical graphs. All data include 3 independent repeats.

### Sencyt correlates with nuclear shape but not cell density in monolayers

Next, we studied the relationship between nuclear shape and Sencyt in cell layers without imposed mechanical perturbations. To this end, we seeded cell monolayers laterally confined by a PDMS gasket. After removing the gasket, cells spread for 24h, leading to monolayer areas with very different cell densities (Figure 3a, f). Performing the same image analyses than in the previous experiments, we observed that the different cell densities also led to different nuclear shapes (Figure 3b, g). Specifically, decreased density, corresponding to increased cell spreading, led to progressive deformation of the nucleus, as indicated by increased oblateness and decreased prolateness (Figure 3b,g, see parameter description in Supp. figure 1). As expected, this also led to an increase in the Solidity index.

**Figure 3.**
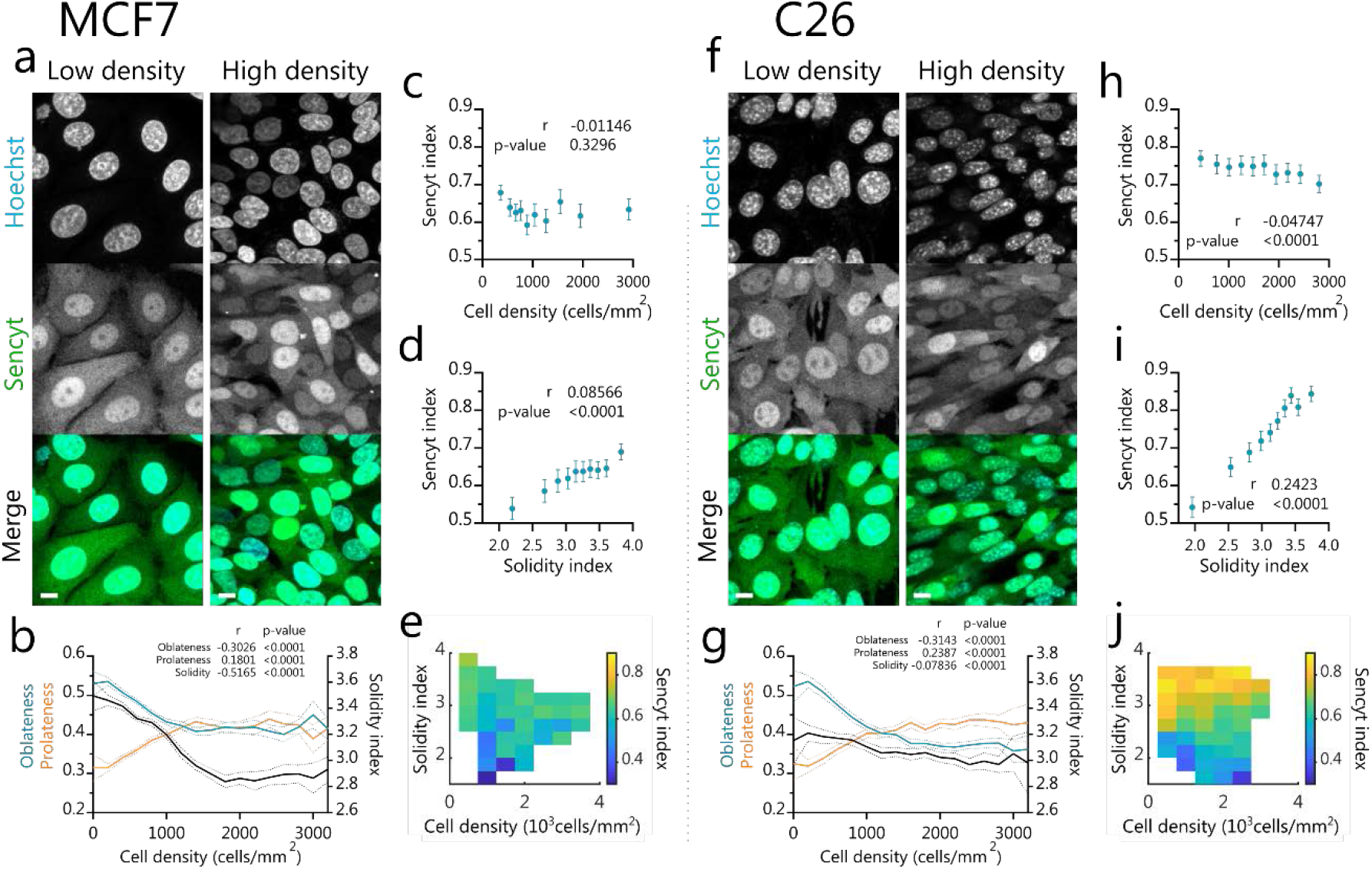
Sencyt correlates better with nuclear shape than cell density in monolayers. a,f) Representative live images of cells in low and high density showing Sencyt and nuclei staining. Scale bars are 10 µm. b,g) nuclear shape parameters versus density (N=20560, 16711 cells), c,h) Sencyt index versus cell density (N=7237, 7522 cells), d,i) Sencyt index versus Solidity index (N=7865, 7647 cells), e,j) Average Sencyt index as a function of Solidity index and cell density, for both cell types (N= 7118, 7522 cells). p-values calculated with Two-tailed non-parametric Spearman correlation test. Error shading and error bars represent 95% CI. All data include 3 independent repeats.

Then we analyzed separately the effect of density and Solidity index in the Sencyt index. The Sencyt index did not correlate with cell density for MCF7 (Figure 3c) and correlated but very mildly for C26 (Figure 3h). In contrast, the Sencyt index correlated with the Solidity index in both cell lines, with a higher correlation for C26, and milder in MCF7 (Figure 3d,i). In summary, Sencyt index correlated much more with the Solidity index than with cell density for both cell lines (Figure 3e, j). In fact, solidity was the nuclear geometrical parameter that best correlated with the Sencyt index for both cell types (Supp. figure 4). Interestingly, C26 cells not only exhibited higher correlations between Sencyt and overall nuclear shape parameters, but also a higher range of Sencyt index values in response to nuclear shape parameters. Differences between cell lines could arise from several factors, including different nuclear mechanical properties, which depend on cell type (Hobson et al., 2020; Kechagia et al., 2023). This may lead to different tension/shape relationships.

### Cell layers show different regulation for Sencyt and for YAP

Finally, we set out to understand if NCT in monolayers is affected in the same way as YAP, a well-known transcription factor that has NCT-regulated mechanosensitivity (Elosegui-Artola et al., 2017), but which also undergoes complex biochemical regulation through the Hippo pathway (Piccolo et al., 2014). To this end, we immunostained for YAP the same cell samples we imaged live for Sencyt (Figure 4a-b, g-h). The YAP nuclear-to-cytoplasmic concentration ratio strongly correlated with density in both cell lines (Figure 4c, i). This is an expected behavior since YAP nuclear localization has been proven to depend on cell-cell contacts: an increase of cell-cell contacts decreases nuclear localization, decreasing proliferation (Aragona et al., 2013; Dupont et al., 2011; Zhao et al., 2007, 2008).

**Figure 4.**
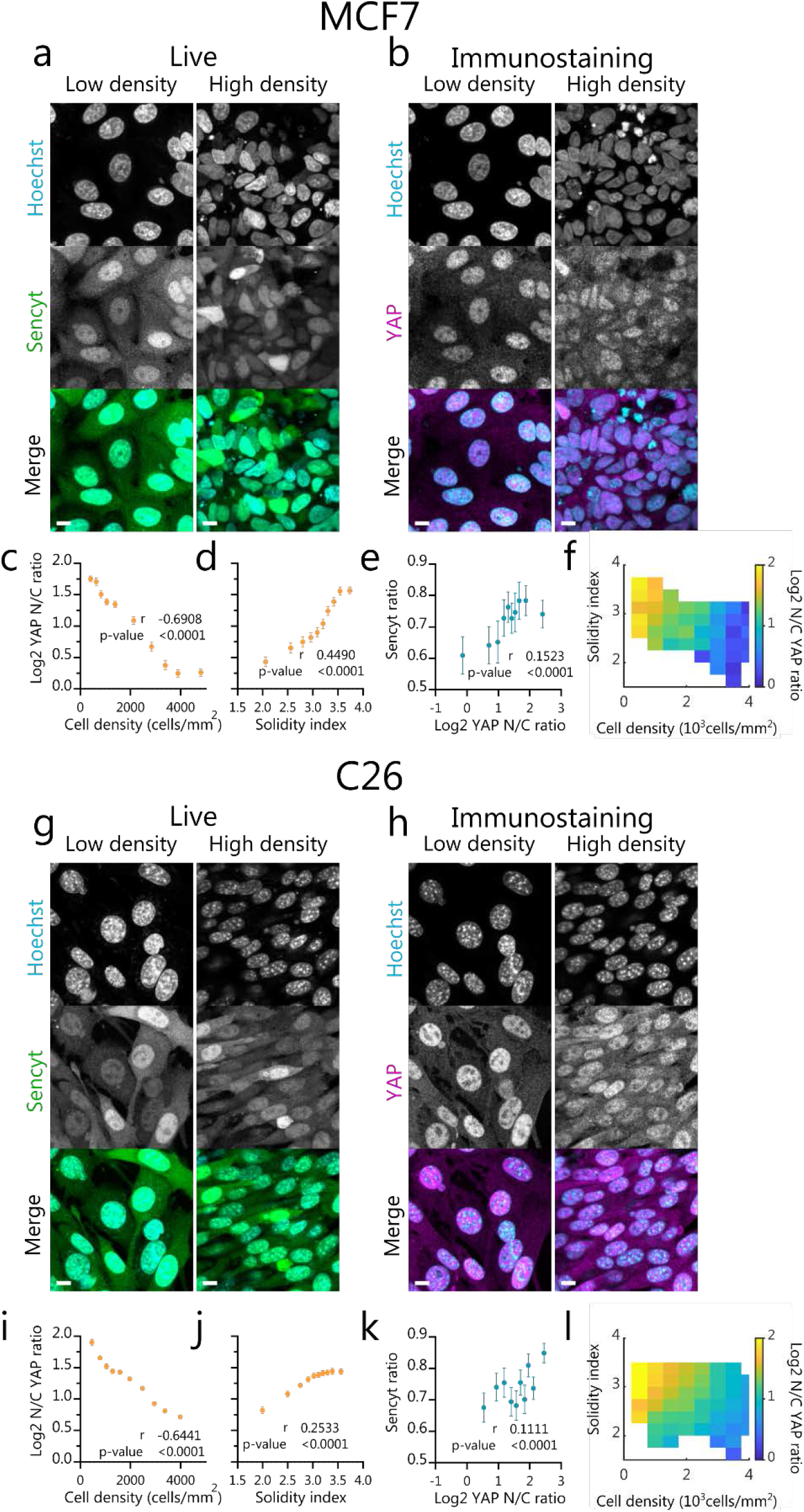
Cell layers show different regulation for Sencyt and for YAP. a,g) Representative images of cells in low and high density showing Sencyt and nuclei staining. b,h) representative images of the fixed same cells in low and high density for YAP immunostaining and nuclei staining. Scale bars are 10 µm. c,i) Log2 N/C YAP ratio versus cell density (N= 4889, 7204 cells), d,j) Log2 N/C YAP ratio versus Solidity index (N= 4889, 7204 cells), e,k) cell-by-cell correlation of Sencyt index versus Log2 N/C YAP ratio (N= 1548, 1782 cells), f,l) Log2 N/C YAP ratio in colour versus cell density and Solidity index, both for MCF7 and C26 cell lines, respectively. (N= 4679, 7050 cells). p-values calculated with Two-tailed non-parametric Spearman correlation test. Error error bars represent 95% CI. All data include 3 independent repeats.

The YAP N/C ratio also correlated with solidity in both cell lines (Figure 4d, j), although to a lesser degree than with cell density. Overall, the combined effects of both factors were clearly visible (Figure 4f, l). Correlations with solidity were similar for YAP and Sencyt in C26 cells (Figure 3I, Figure 4j), but much higher for YAP in MCF7 cells (Figure 3d, 4d). Further, for both cell lines Sencyt and YAP localization correlated significantly with each other, more in MCF-7 than in C-26 (Figure 4e, k). However, correlations were low, likely reflecting the fact that both parameters are not molecularly tied, and that the different layers of YAP regulation reduce the correlations. In fact, YAP localization correlated better than Sencyt index with most nuclear shape and other geometrical parameters (Supp. figure 4).

## Discussion

In this work, we harness the Sencyt sensor to demonstrate that NCT in cell monolayers is regulated by mechanical stimuli, leading to altered nuclear accumulation of shuttling proteins. The role of mechanics in NCT had been previously demonstrated in single cells, in response to either increased substrate stiffness, force applied to the nucleus (Andreu et al., 2022; Elosegui-Artola et al., 2017), or hypo-osmotic shocks (Zimmerli et al., 2021). Here, we demonstrate that mechanics also plays a role in multicellular systems, in response to both hypo and hyper-osmotic shocks, and to cell contractility. The mechanism involved is likely the same: increased nuclear membrane tension caused by applied force (Dahl et al., 2004; Enyedi et al., 2016), a subsequent increase in the diameter of the NPC (Zimmerli et al., 2021), and resulting differential alteration in passive versus facilitated diffusion through NPCs (Andreu et al., 2022).

Certainly, other factors beyond nuclear envelope tension could also be playing a role. Hyper-osmotic shocks have been shown to slow intracellular signaling due to molecular crowding (Miermont et al., 2013), and to decrease nuclear import (Ng et al., 2014) by impairing the Ran system (Kelley & Paschal, 2007). Further, osmotic swelling due to tissue damage, and subsequently increased nuclear envelope tension, induces signaling by translocating cytosolic phospholipase A2 (cPLA2) and 5-lipoxygenase (5-LOX) into the inner nuclear membrane (Cho & Stahelin, 2005; Enyedi et al., 2016). The contribution of these different mechanisms in response to specific perturbations remains to be elucidated. However, the common response to very different stimuli (osmotic shocks versus contractility inhibition), combined with the correlation with nuclear solidity, strongly suggest a role of nuclear envelope tension, and direct effects on NPC permeability.

Comparing some of our results leads to interesting implications. First, mechanically induced changes in NCT can occur both with nuclear deformation (in response to osmotic shocks) and without (in response to contractility inhibition). This suggests that mechanical perturbations can affect NCT and likely nuclear envelope tension without visible changes in nuclear shape. This could potentially be explained by different means of transmitting force: through global nuclear swelling in the case of osmotic shocks, versus specifically through the LINC complex in response to cell contractility. Indeed, in our previous work we found a very different response of Sencyt in single cells when nuclei were deformed in the presence or absence of LINC complexes (Andreu et al., 2022). Such differences could also explain the fact that nuclear solidity and the Sencyt index correlate, but with rather low correlation values. Still, larger deformations will lead to larger effects, as indicated by the correlations between Sencyt index and Solidity index.

Second, we found very interesting differences when comparing Sencyt versus YAP responses. Importantly, both Sencyt and YAP responded to nuclear solidity, but only YAP showed a clear response to cell density. This differential behavior allowed us to decouple the effects of mechanics and cell-cell adhesion, showing that the role of cell-cell adhesion in YAP cannot be explained by mechanical effects on NCT. This is likely due to the several layers of YAP regulation, and specifically the role of cell-cell adhesion in the hippo pathway (Aragona et al., 2013; Zhao et al., 2007, 2008). Interestingly, even if it also responded to density, YAP correlated better than Sencyt with nuclear solidity, at least for MCF7 cells. This suggests that YAP properties may have evolved to result in a more optimal mechanosensor than the synthetic Sencyt. In the future, further optimization of Sencyt may increase sensitivity, potentially revealing interesting insights on how physiological mechanosensitive molecules such as YAP evolved.

Methodologically, our results also show that a sensor of nucleocytoplasmic transport (Sencyt), together with image analysis, is a valuable tool to understand NCT regulation in multicellular environments *in vitro* and potentially *in vivo*, merely by using confocal fluorescence in live imaging. Using Sencyt is likely to reveal much finer NCT regulation than merely employing fluorophores tagged with a strong NLS sequence, as those strongly localize to the nucleus unless NCT is acutely disrupted. Potentially, Sencyt may for instance be used to identify other mechanosensitive transcriptional regulators purely regulated by NCT and not cell-cell adhesion in multicellular systems, overriding YAP-like regulation systems. Beyond mechanics, it could also be used to study alterations in NCT due to any other factor.

## Methods

### Cell lines

MCF-7 and C-26 cells were cultured in Dulbecco’s modified Eagle medium (DMEM) supplemented with fetal bovine serum (FBS, 10% aq.), L-glutamine (2 mM), penicillin (100 U/mL) and streptomycin (100 µg/mL) in a humidified atmosphere with 5% CO2 at 37 °C in humidified atmosphere. C-26 was kindly offered by Onno Kranenburg. A plasmid transiently expressing Sencyt was previously described as L_NLS 41kDa (Andreu et al., 2022) and is available in addgene (Addgene plasmid # 201342 ; http://n2t.net/addgene:201342 ; RRID: Addgene_201342). For the creation of stable cell lines expressing Sencyt, pLentiPGK coding for SV40A4-EGFP-2PrA (Andreu et al., 2022) was cloned using these primers to excise it from the parental plasmid: Infusion_SV40A4-EGFP-2PrA_Fwd cgg tac cgc ggg ccc atg ggc cca aaa aag gc; Infusion_SV40A4-EGFP-2PrA_Rev gaa agc tgg gtc tag acc act ttg tac aag aaa gct ggg tcg g. The plasmid was then used for viral production in HEK293T (ATCC^®^ CRL-1573™) of low passage in media IMDM supplemented with 10% heat-inactivated FCS, 1% pen/strep. Reagents used were: 2.5 M CaCl2, 0.1x TE buffer, 2x HBS pH 7.12 (fresh). Cell lines were transduced with a mix of supernatant containing virus and polybrene (Sigma H9268 suspended at 4mg/ml in sterile water, 1:1000), at 37°C for 24 hours. Transduced cells were selected by Hygromycin 200µg/ml, and a FACS sorting procedure based on GFP fluorescence.

### Transient transfection

Cells were transfected the day before the experiment using Neon transfection device (Thermo Fisher Scientific) according to the manufacturer’s instructions. MCF-7: Pulse voltage 1250 V; Pulse width 20 ms; Pulse Number 2. C-26: Pulse voltage 1350 V; Pulse width 20 ms; Pulse Number 2. pcDNA3.1-mCherry was a gift from David Bartel (Addgene plasmid # 128744; http://n2t.net/addgene:128744;RRID:Addgene_128744).

### Imaging settings

Image acquisition was done with a Zeiss LSM880 inverted confocal microscope objective and using Zeiss ZEN2.3 SP1 FP3 (black, version 14.0.24.201), using a 63X 1.46 NA oil immersion objective and a 403, 488, 561 and 633nm wavelength lasers, in the Fast Airyscan mode. Voxel size was of 0.1413 µm for xy and z-step of 0.4 µm. This allowed us to activate the Definite Focus system, so the sample was autofocused every time frame. Pixel sizes are of 0.1409 µm, and z-spacing for the objective is 0.4 microns, which turns into 0.3440 µm after correction. Z-spacing was corrected following the literature (Diel et al., 2020), considering cell refractive index of 1.36 and Immersol immersion oil of 1.518.

For cell layers image positioning was automatically set to fit a tile positioning with an 15% image overlap. In the case of YAP immunostaining for cell layers, only properly permeabilized regions were imaged. To recognize the properly permeabilized regions a control staining of Sencyt was performed (not shown).

### Osmotic shock experiments

#### Cell seeding

Single-well, Mattek, glass-bottom dishes were incubated with 10 μg/mL of fibronectin in PBS for 1 hour at room temperature. Cells were seeded on the plate to achieve an approximate density of 1000 cells/mm^2^ the day after. Minimum 1h prior to experiment, the medium of the cells was changed to 500µL of medium containing 1/10.000 Hoechst 33342 (Invitrogen).

#### Image acquisition and Osmotic shock

The time frame was set to 30s, for every sample we imaged 5 timepoints without disturbances before the shock was applied. Then we imaged for 45 timepoints more. To decrease image drift while acquiring images of the same position through time, we started the imaging of the sample with 0.5mL of medium containing the nuclei stain. At the time of the shock, we added 1mL of 1.5x solution either for hypo or hyper-osmotic shock conditions. The control worked as an imaging control condition.

Cell medium has an osmolarity of ∼340 mOsm. ∼113 mOsm hypo-osmotic shocks (66%) were performed by mixing the 500µL of medium with 1.5x de-ionized water with Ca2+ and Mg2+ ion concentration corrected to match those of the medium (264mg/L CaCl2 · 2H2O, 164.67mg/L MgCl2 · 6H2O). ∼695 mOsm hyper-osmotic shocks (204%) were performed by adding 1mL of 1.5x solution containing 96.9g/L D-mannitol (Sigma) to the medium.

For the analyses, t=30 s was discarded because it was noisy due to out of focus imaging after the medium pippeting.

### Drug treatment experiments

#### Cell seeding

6-well, Mattek, glass-bottom dishes were incubated with 10 μg/mL of fibronectin in PBS for 2 hours at room temperature. Cells were seeded on the plate in mediums containing 1/10.000 Hoechst 33342 (Invitrogen) and either the drug vehicle (3.5 µL DMSO/1 mL of medium) or a combination of the drugs (25 µM para-NitroBlebbistatin, 50 µM CK666). Cells were left to attach to the substrate for 2 hours before the imaging started.

#### Image Acquisition and Drug washout

Image acquisition parameters were identical as the Osmotic Shock experiments unless specified otherwise. For the Drug washout experiment, cells were imaged every hour, starting 2 h after seeding. For 2 timepoints cells were left untouched. After the imaging of the 2 timepoint finished we aspirated the drug-containing medium of the condition of the Drug washout, washed twice with warm medium, and added the vehicle containing medium for the following timepoints.

### Cell layer experiments

#### Cell seeding

Mattek, glass-bottom dishes were incubated with 10 μg/mL of fibronectin in PBS for 2 hours at room temperature. Magnetic PDMS gaskets (Rodriguez-Franco et al., 2017) sized 4mm times 8mm at the inner side, were treated water and soap, washed in EtOH, washed in MiliQ, incubated in Pluronic^®^ F-127 (20g/L) 1h room temperature, washed twice in PBS, and air dried. Both Matteks and gaskets were UV sterilized before seeding. For cell seeding, gaskets were put in the center of the Matteks dishes, and the dishes were placed on top of a holder including a magnet to keep them in place. Approximately 60k cells were seeded in every gasket (0.3 cm^2^). Cells were incubated for 4h, and then some washes with medium were performed to retrieve non-attached cells. Enough medium was added to cover the gaskets completely. Cells were then incubated for 24h with the gasket. The gasket was then retrieved, and cells were incubated O/N before imaging started.

#### Staining

Immunostainings were performed as previously described (Elosegui-Artola et al., 2017). Cells were fixed with 4% v/v paraformaldehyde for 10 minutes, permeabilized and blocked with 0.1% (MCF-7) and 1% (C-26) v/v Triton X-100 and with 2% v/v Fish-Gelatin in PBS 1X for 45 minutes, incubated with primary antibody for 1 hour at room temperature or O/N at 4ºC, washed 3 times with Fish-Gelatin-PBS for 5 minutes, incubated with secondary antibody for 1 hour, washed with Fish-Gelatin-PBS 3X for 5 minutes, and image in PBS with the same conditions as the live imaging. YAP mouse monoclonal antibody (Cat# sc101199; RRID: AB_1131430), and secondary Alexa Fluor-555 (ThermoFisher, goat anti-Mouse, A-21424; RRID: AB_141780) were used diluted 1:400.

### Image analysis

Images were processed to .czi format with Zeiss ZEN2.3 SP1 FP3 (black, version 14.0.24.201). Then they were binned in xy by a factor of 4 calculating the median using Fiji (Schindelin et al., 2012), leaving the voxel size in xy at 0.5652 µm (z remained untouched, values were averaged) then they were separated by channels (the nuclei staining was filtered with a median filter of 2 pixels, for osmotic shock experiments, to decrease the effects of chromatin staining changes into segmentation). The processed nuclei image was then segmented in 3D using Cellpose (Stringer et al., 2021).

Code used for Osmotic shocks experiments: *python -m cellpose --dir* [directory] *--do_3D -- cellprob_threshold=-2*.*0 --batch_size 2 --pretrained_model nuclei --chan 1 --diameter 34. -- save_tif --no_npy --use_gpu --verbose --anisotropy 0*.*6*.

Code used for Drug washout experiments: *python -m cellpose --dir* [directory] *--do_3D -- cellprob_threshold=-2*.*0 --batch_size 2 --pretrained_model nuclei --chan 1 --diameter 34. -- save_tif --no_npy --use_gpu --verbose*

Code used for cell layer experiments: *python -m cellpose --dir* [directory] *--do_3D -- cellprob_threshold=0*.*0 --batch_size 2 --pretrained_model nuclei --chan 1 -- diameter 34. -- save_tif --no_npy --use_gpu --verbose*

Using the masks created by Cellpose we measured fluorescent intensities inside and outside of the nucleus for the plane of biggest area for every nucleus. The nuclear area was created by eroding this plane by 1 pixel (0.5652 µm), and the cytoplasmic by creating a ring outside the nucleus. This was done by subtracting a 3-pixel-increased area, by a 1-pixel-increased area. Then any pixel in the cytoplasmic area was excluded if it fell inside any neighbouring nucleus. This was done for all channels, as well as measuring geometrical and size parameters of the masks using MATLAB2020b. To avoid spurious measurements some filters were applied. For cell brightness: minical signal to noise ratio filter was applied. For dim cells next to very bright cell, or vice versa: all measurements of areas with a coefficient of variation higher than 0.8 were discarded. For bad nuclei segmentation: nuclei with nuclear-to-cytoplasmic brightness ratios of Hoechst lower than 4 were discarded.

### Data quantification and parameters

Once the nuclei masks were obtained, in MATLAB we calculated the different nuclear shape parameters (see Supp. figure 1) in two ways: directly from the mask (Volume, Solidity index) and fitting an ellipsoid and obtaining the length of the three radii (Oblateness, Prolateness). In the case of the Solidity index, the convex hull volume is the smallest convex volume that contains a shape.

Sencyt index was calculated as the logarithm in base 2 of the ratio of the mean nuclear fluorescence 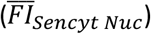 and the mean cytoplasmic fluorescence 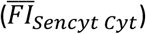 of Sencyt, after subtracting mean background fluorescence 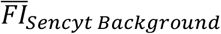, assessed in cell-free regions of the image):

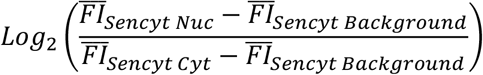

Log2 N/C YAP ratio was calculated as the logarithm in base 2 of the ratio of the mean nuclear fluorescence 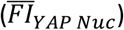 and the mean cytoplasmic fluorescence 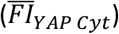 of YAP staining, also after subtracting mean background fluorescence 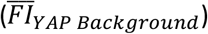:

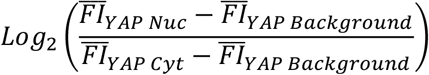

For Solidity index, Sphericity, Oblateness and Prolateness, see Supp. figure 1.

Cell Density was calculated with the xy coordinates by measuring the number of nuclei around the n^th^ nucleus. This was done by centering a square in the nth nucleus with a side size of 200 pixels, which is 113.03 µm.

For Figure 4e, k, direct nuclei correlation between the live Sencyt images and the YAP staining images was done by nuclei image registration and overlap of the masks. This way we obtained a table with the corresponding mask identifiers in live and staining images.

## Acknowledgements

We thank S. Usieto, A. Menédez, N. Castro, M. Purciolas-Casas and M. Morcillo for providing technical support. We acknowledge funding from “la Caixa” Foundation (grant LCF/PR/HR20/52400004 to X.T., J.V.R., and P.R.-C. and fellowships LCF/BQ/DR19/11740009 to I.G.-M. and LCF/BQ/DR19/11740009 to M.M.J.), the Spanish Ministry of Science and Innovation (PID2022-142672NB-I00 to P.R.-C., PID2021-128635NB-I00 to X.T., and PID2022-141040NA-I00 to I.A.), the Generalitat de Catalunya (2021 SGR 01425 to X.T. and P.R.-C.), The prize “ICREA Academia” for excellence in research to P.R.-C., Fundació la Marató de TV3 (201936-30-31 to P.R.-C.), the European Research Council (grant 101097753 MechanoSynth to P.R-C. and Adv-883739 Epifold to X.T.), and The Basque government (grant PIBA_2023_1_0033 to I.A.). This work was supported in part by the Fundación Biofísica Bizkaia and the Basque Excellence Research Centre (BERC) program of the Basque Government. IBEC is a recipient of a Severo Ochoa Award of Excellence from MINCIN.

## Author contributions

Conceptualization: I.G.-M., I.A., P.R.-C.

Resources: I.G.-M., G.B., X.T., J.V.R., I.A.

Data curation: I.G.-M., I.A.

Software: I.G.-M.

Formal analysis: I.G.-M., I.A.

Supervision: I.A., P.R.-C.

Funding acquisition: X.T., J.V.R., I.A., P.R.-C.

Validation: I.A., P.R.-C.

Investigation: I.G.-M., B.G., M.M., M.G., I.A.

Visualization: I.G.-M.

Methodology: I.G.-M., G.B., X.T., J.V.R., I.A.

Writing—original draft: I.G.-M.

Project administration: P.R.-C.

Writing—review and editing: I.G.-M., I.A., P.R.-C.

## Data availability

Source data for all figures is available as a supplementary file.

## Figures

**Supp. figure 1.**
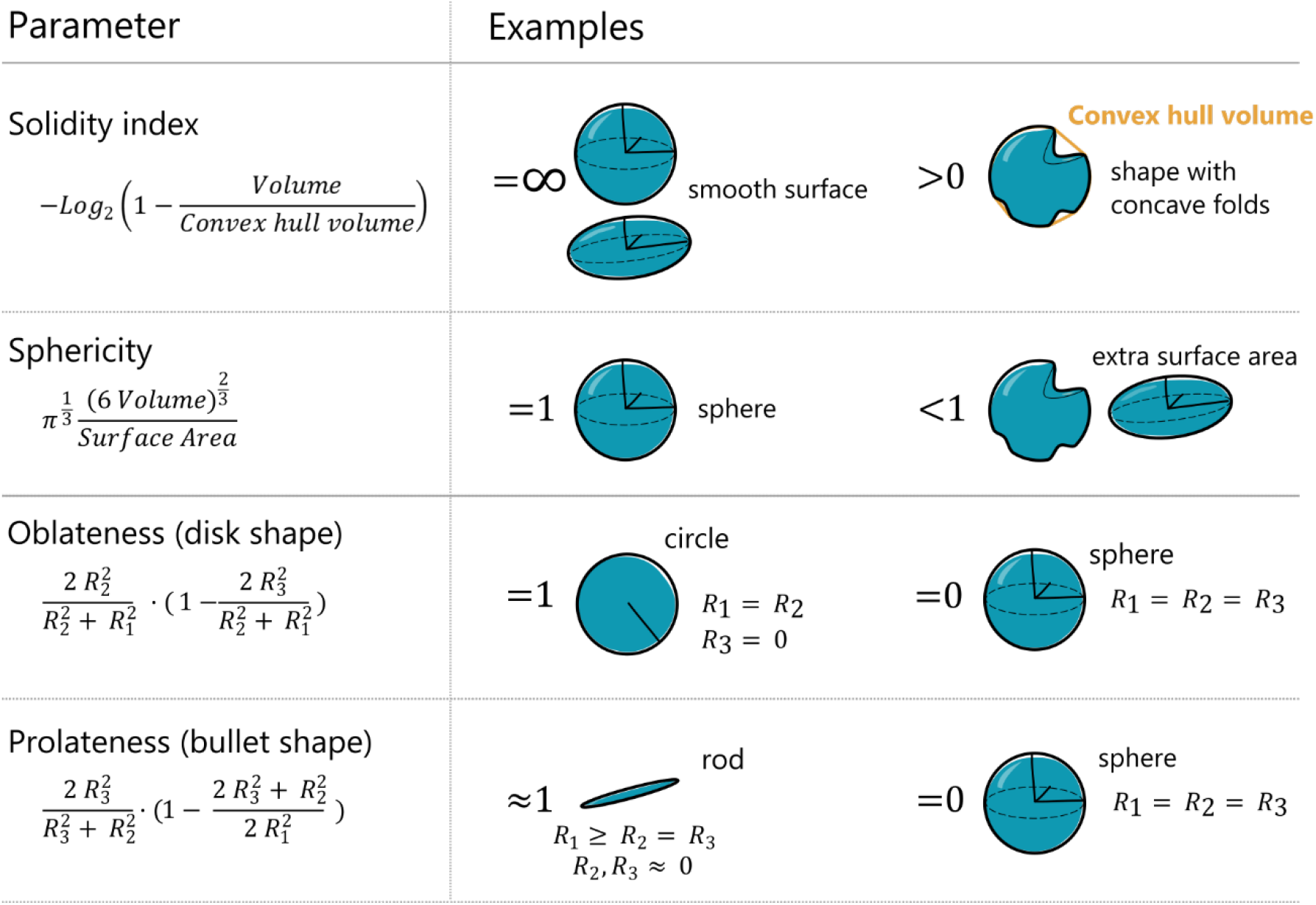
Nuclear shape parameters description. R_1_, R_2_, R_3_ correspond to the three radii (from the largest to the smallest; R_1 ≥_ R_2 ≥_ R_3_) of an ellipsoid fitted to the segmented nucleus. Volume and surface areas are the measured volume and surface area of the nucleus as obtained from nuclear segmentation. Convex hull volume is the volume of the convex hull, defined as the smallest convex shape (that is, not containing any concave folds) that encloses the nucleus.

**Supp. figure 2.**
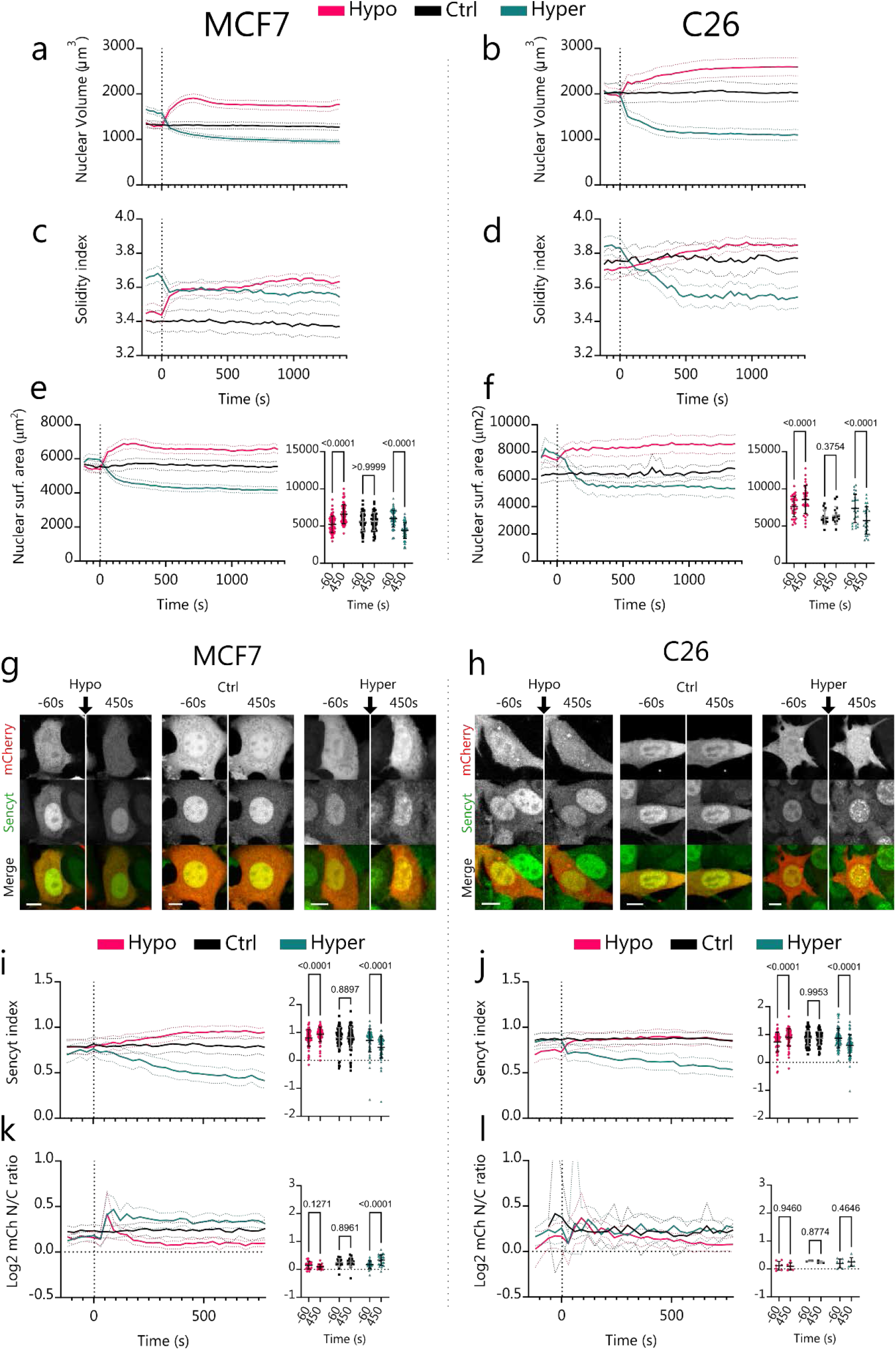
Additional measurements of effects of osmotic shocks. Absolute values for Nuclear Volume (a-b) (N=170, 130, 188, 81, 72, 107 cells) and Solidity index (c-d) (N=230, 170, 231, 121, 104, 142 cells), for MCF7 an,, d C26, respectively. e-f) Change of nuclear surface area over time for MCF7 and C26, with corresponding statistics (N=68, 50, 59, 32, 15, 26 cells). g-h) Representative images of cells transfected with mCherry, submitted to osmotic shocks as in Figure 1. Scale bar is 10 µm. i-j) Corresponding quantification of Sencyt index (N=71, 78, 73, 58, 67, 69 cells) and k-l) Log2 mCherry Nucleo-cytoplasmic ratio (N=20, 20, 27, 7, 3, 7 cells). p-values calculated with 2-way ANOVA corrected with Šídák’s multiple comparisons test. Error bars represent 95% CI for timelapse graphs and SD for statistical graphs. All data include 3 independent repeats.

**Supp. figure 3.**
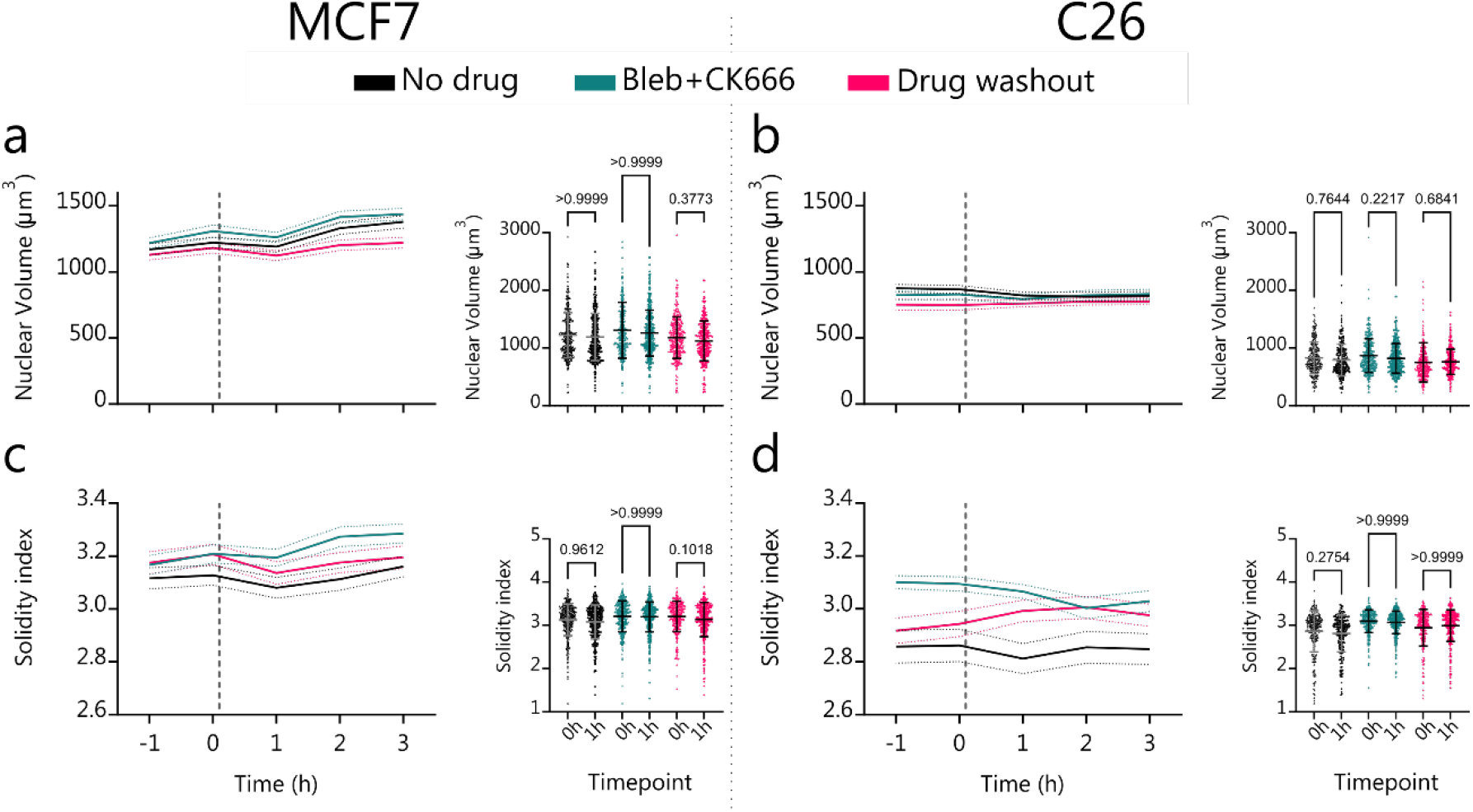
Nuclear volume and Solidity index values corresponding to Figure 2 data. a-b) Nuclear Volume (N=368, 370, 398, 436, 335, 332, 230, 218, 368, 394, 303, 302 cells) and c-d) Solidity index measurements and statistics for MCF7 and C26 as a function of time. (N=368, 370, 398, 436, 335, 332, 230, 218, 368, 394, 303, 302 cells). p-values calculated with Kruskal-Wallis test corrected with Dunn’s multiple comparisons test. Error bars represent 95% CI for timelapse graphs and SD for statistical graphs. All data include 3 independent repeats.

**Supp. figure 4.**
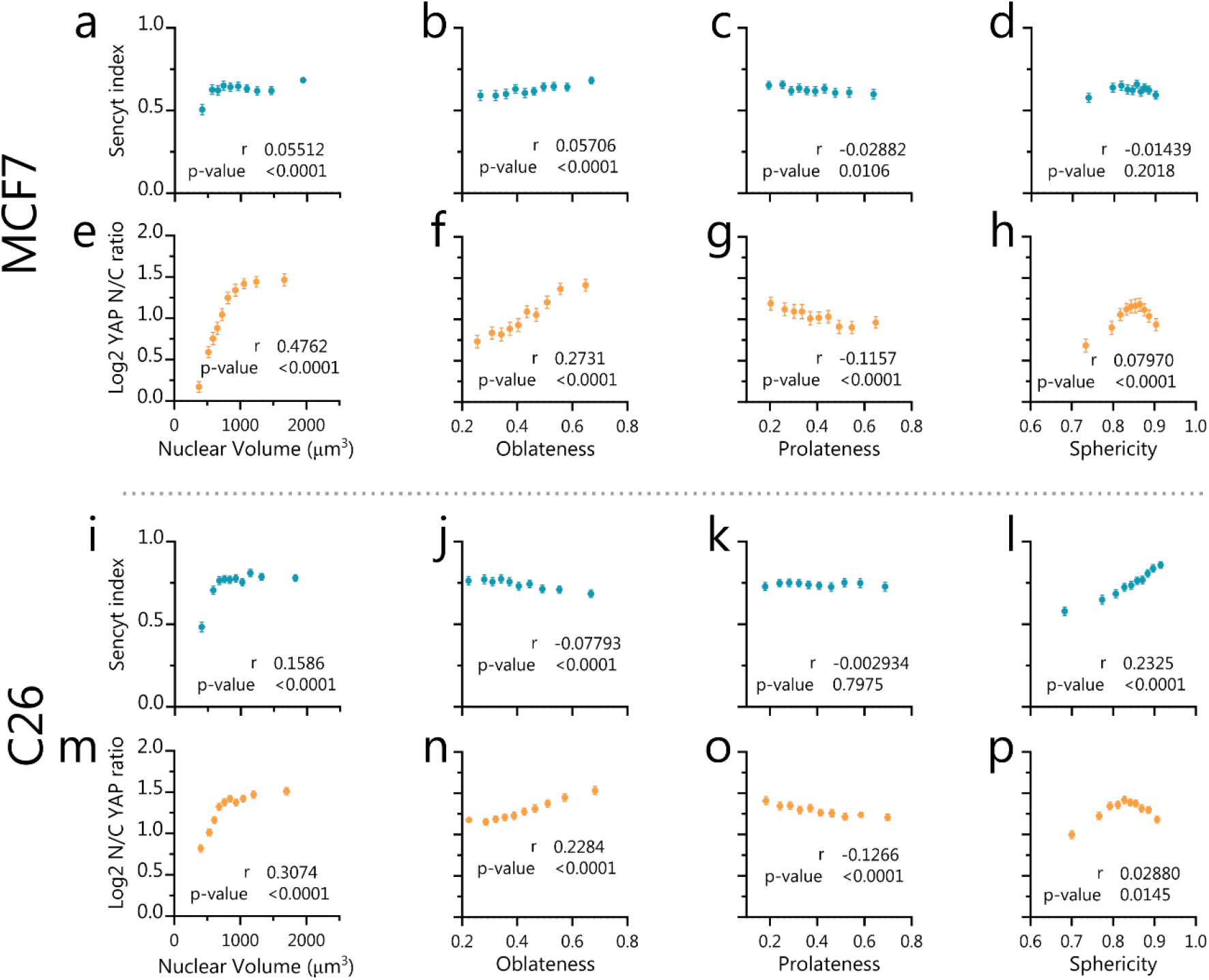
Sencyt index and YAP localization versus nuclear shape parameters. (a-d N=7865 cells, e-h N=4889 cells, i-l N=7647 cells, m-p N=7204 cells). p-values calculated with Two-tailed non-parametric Spearman correlation test. Error error bars represent 95% CI. All data include 3 independent repeats.

